# Arrhythmia-associated Calmodulin Variants Interact with KCNQ1 to Confer Aberrant Membrane Trafficking and Function

**DOI:** 10.1101/2023.01.28.526031

**Authors:** Po wei Kang, Lucy Woodbury, Paweorn Angsutararux, Namit Sambare, Jingyi Shi, Martina Marras, Carlota Abella, Anish Bedi, DeShawn Zinn, Jianmin Cui, Jonathan R. Silva

## Abstract

**Rationale:** Missense variants in calmodulin (CaM) predispose patients to arrhythmias associated with high mortality rates. As CaM regulates several key cardiac ion channels, a mechanistic understanding of CaM variant-associated arrhythmias requires elucidating individual CaM variant effect on distinct channels. One key CaM regulatory target is the KCNQ1 (K_V_7.1) voltage-gated potassium channel that underlie the I_Ks_ current. Yet, relatively little is known as to how CaM variants interact with KCNQ1 or affect its function.

**Objective:** To observe how arrhythmia-associated CaM variants affect binding to KCNQ1, channel membrane trafficking, and KCNQ1 function.

**Methods and Results:** We combine a live-cell FRET binding assay, fluorescence trafficking assay, and functional electrophysiology to characterize >10 arrhythmia-associated CaM variants effect on KCNQ1. We identify one variant (G114W) that exhibits severely weakened binding to KCNQ1 but find that most other CaM variants interact with similar binding affinity to KCNQ1 when compared to CaM wild-type over physiological Ca^2+^ ranges. We further identify several CaM variants that affect KCNQ1 and I_Ks_ membrane trafficking and/or baseline current activation kinetics, thereby contextualizing KCNQ1 dysfunction in calmodulinopathy. Lastly, we delineate CaM variants with no effect on KCNQ1 function.

**Conclusions:** This study provides comprehensive functional data that reveal how CaM variants contribute to creating a pro-arrhythmic substrate by causing abnormal KCNQ1 membrane trafficking and current conduction. We find that CaM variant regulation of KCNQ1 is not uniform with effects varying from benign to significant loss of function. This study provides a new approach to collecting details of CaM binding that are key for understanding how CaM variants predispose patients to arrhythmia via the dysregulation of multiple cardiac ion channels.

## Introduction

Calmodulin (CaM) is a ubiquitous auxiliary subunit of key ion channels underlying the cardiac action potential. In the heart, CaM regulatory targets include ryanodine receptors (RyR2) and several voltage-gated channels including calcium channels (e.g. CaV1.2), sodium channels (e.g. NaV1.5), and potassium channels (e.g. KCNQ1 or K_V_7.1) ^1-6^. CaM canonically acts as a Ca^2+^ sensor and confers Ca^2+^ sensitivity to channels to allow Ca^2+^-dependent inactivation in CaV1.2 and NaV1.4 ^1,3,7^. Independent of Ca^2+^-sensing function, CaM binding also modulates baseline channel function in CaV1.2, NaV1.5, and KCNQ1 ^8-11^. Inherited or *de novo* mutations in these channels causing CaM dysregulation are associated with arrhythmias such as Timothy syndrome (TS), Brugada syndrome (BrS), and long QT syndrome (LQTS) ^9,12-14^. Three genes in the human genome (*CALM1-3*) encode for CaM with 100% conservation in amino acid identity ^3,15^, and CaM was long-thought to be intolerant of primary sequence alterations. Recently, human missense variants in CaM have emerged as a molecular factor underlying arrhythmias (“calmodulinopathy”) such as LQTS and catecholaminergic polymorphic ventricular tachycardia (CPVT) associated with high mortality rate ^14,16-20^. As CaM variants cause dysfunction in many combinations of their regulatory targets, calmodulinopathy mechanisms are fundamentally complex. Extensive studies have established that several CaM missense variants impair Ca^2+^ binding ^16,21-28^. To date, CaM variants are most extensively evaluated as related to CaV1.2 or RyR2, with many variants shown to induce CaV1.2 and RyR2 dysfunction thereby contributing to arrhythmogenesis ^14,22,27,29-34^. By comparison, relatively little is known regarding whether CaM variants may also alter KCNQ1 channel function and membrane expression, with only a few studies suggesting potential effects ^10,28,35^. Moreover, the relative contributions of trafficking, gating effects, and binding are not well-delineated for different variants. These knowledge gaps represent a missing piece for in-depth understanding of calmodulinopathy mechanisms that integrate distinct CaM regulatory targets.

In the heart, KCNQ1 is a homo-tetrameric potassium channel that associates with the auxiliary subunits KCNE1 and CaM to conduct the slow delayed rectifier current (I_Ks_), which participates in action potential repolarization primarily in the context of β-adrenergic stimulation ^5,6,36-38^. Structurally, CaM interacts with Helix A and Helix B within the KCNQ1 carboxy-terminus domain (CTD) as well as the S2-S3 linker within the KCNQ1 voltage-sensing domain (VSD) (Fig. 1A-B) ^5,6,10,35,38,39^. CaM association with KCNQ1 is required for channel assembly and membrane trafficking ^5,6^. CaM further exerts functional effects by triggering current enhancement upon elevated intracellular Ca^2+^ and modulating KCNQ1 and I_Ks_ baseline current under resting Ca^2+^ conditions ^10,39-41^. Mechanistically, KCNQ1 features a gating mechanism leveraged by CaM and KCNE1 to modulate the channel. Upon membrane depolarization, the KCNQ1 VSD activates in a two-step manner, from the resting state to a stable intermediate state, then to the activated state ^38,42-44^. VSD occupancy of the intermediate and the activated states both trigger KCNQ1 pore opening, yielding two open states ^42-44^. KCNE1 suppresses intermediate state conductance to help delay current activation kinetics ^43,44^. CaM controls channel occupancy of the activated state to modulate KCNQ1 current activation kinetics ^10^. Taken together, CaM missense variants have the potential to alter multiple KCNQ1 parameters from trafficking to current conductance, all of which may lead to abnormal action potential repolarization.

**Figure 1.**
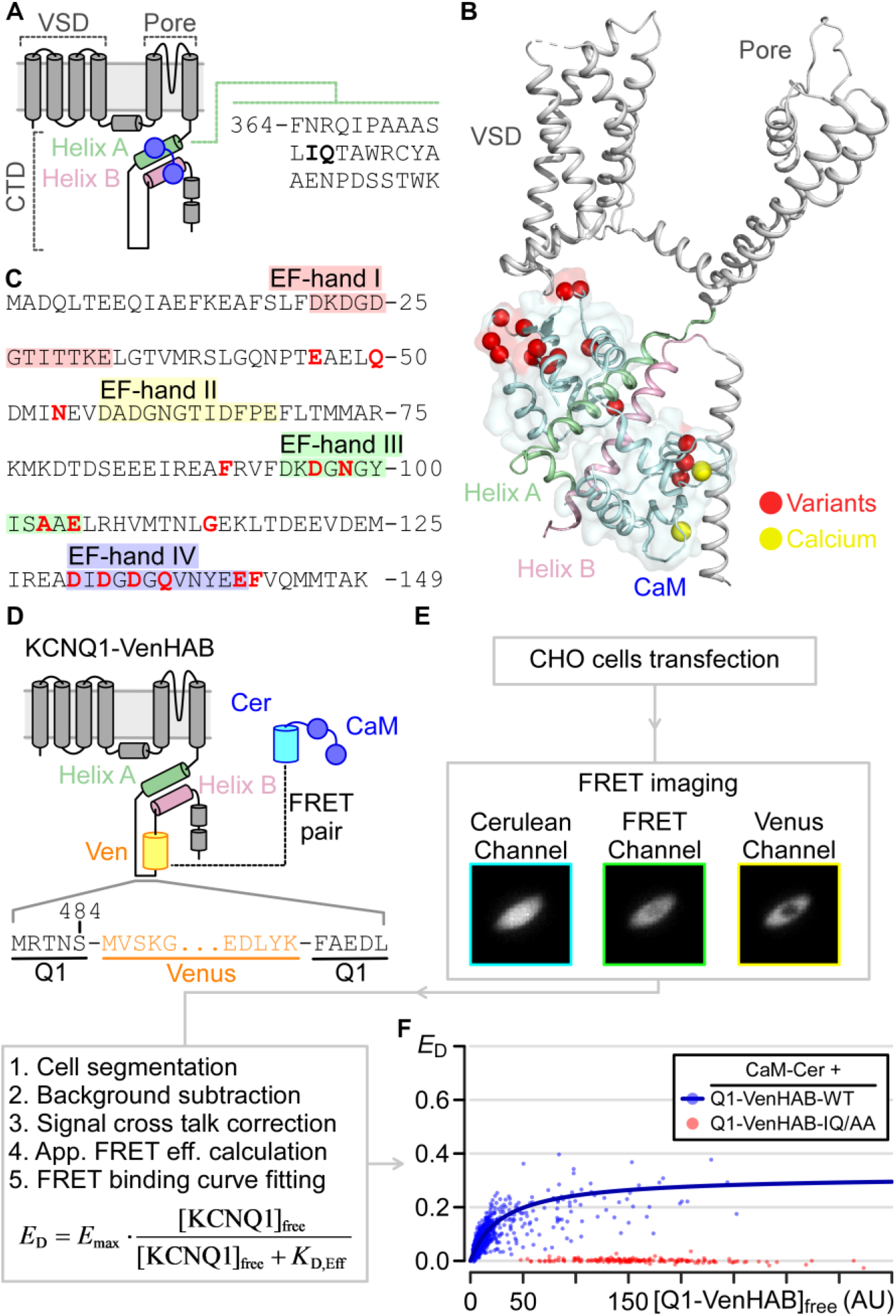
A FRET-based live-cell assay to probe CaM interaction with full-length KCNQ1. **(A)** KCNQ1 topology and CaM interacting regions. CaM binds Helix A (HA) and Helix B (HB) within the C-terminus domain (CTD) of KCNQ1. VSD is voltage-sensing domain. The sequence for KCNQ1 helix A is shown, with the key CaM-interacting “IQ” residues bolded. **(B)** Cryo-EM structural depiction (PDB: 6UZZ) of the human KCNQ1-CaM complex with red spheres indicating positions with known CaM variants. Yellow spheres are calcium ions. **(C)** Sequence of CaM. Each EF hand (colored highlights) can coordinate one Ca^2+^ ion. Red residues correspond to positions with known CaM variants. **(D)** Cartoon illustration of the KCNQ1-VenHAB and CaM-Cerulean FRET pair. The venus insertion site is shown. **(E)** FRET workflow to detect CaM interaction with full-length KCNQ1 in live cells. **(F)** Fitted FRET binding curves of cerulean-tagged CaM with KCNQ1-VenHAB-WT (blue, *n* = 1768) and KCNQ1-VenHAB-IQ/AA (red, n = 217). Each dot is one cell. For WT, best-fit *E*_Max_ = 0.317 and K_D,Eff_ = 30.2 AU with 95% CI = [23.9, 36.7]. For IQ/AA, no fitting was performed because of the lack of rise in FRET signal.

Here, we perform extensive characterization of >10 CaM variants to determine whether KCNQ1 dysfunction contributes to calmodulinopathy-related arrhythmias (Fig. 1B-C). The CaM variants investigated here cover nearly all currently known variant positions. We employ a novel fluorescence resonance energy transfer (FRET) assay to quantify CaM variants binding to full-length KCNQ1 in live cells, thereby identifying CaM variants capable of competing with CaM wild-type (WT) for binding KCNQ1. Using fluorescence assays and functional electrophysiology, we further identify specific CaM variants that confer abnormal KCNQ1 membrane trafficking or aberrant current conduction. In all, our results systematically map distinct CaM variant effects to KCNQ1 binding, membrane trafficking, and current conduction, delineating how KCNQ1 dysfunction contributes to calmodulinopathy.

## Results

### A FRET-based assay to probe CaM interaction with full-length KCNQ1

Although calmodulin is encoded by three genes in the human genome, the presence of one gene carrying a missense variant is sufficient to induce arrhythmia ^16-19^. This genetic pattern suggests that CaM variants and CaM WT are both expressed in cardiomyocytes. Thus, CaM variants competes with CaM WT for binding to their regulatory targets. As KCNQ1 channel assembly and membrane expression require CaM association ^5,6^, the ability for CaM variants to bind KCNQ1 figures critically into whether the variant may impart current dysfunction on membrane-expressing KCNQ1. To explore this question, we developed a FRET assay to quantify binding between CaM and KCNQ1 channels in live cells. FRET is a phenomenon in which nanometer proximity (< ∼100Å) between two appropriate fluorophores can be optically detected and has been successfully employed to study CaM-channel interaction ^29,45-48^. Here, we innovated on prior techniques through the design of a novel FRET pairs between CaM and full-length KCNQ1, as well as the development of an EMCCD camera-based imaging technique and automated analysis to efficiently quantify binding between CaM variants and KCNQ1 (see extended methods).

In developing the FRET assay, we first selected the well-characterized Cerulean-Venus FRET pair ^49^ and screened for optimal fluorophore placement to detect FRET signal between CaM and the full-length KCNQ1. Cerulean was labeled to the CaM carboxy-terminus by fusion, enabling the Cerulean-tagged CaM to act as a FRET donor (Fig. 1D). For the FRET acceptor, we screened for optimal labeling position by fusing a Venus to the carboxy-terminus or the cytosolic Helix A – Helix B (HAB) linker of the full-length KCNQ1. Insertion of a Venus fluorophore within the KCNQ1 HAB linker after residue S484, which we term KCNQ1-VenHAB, yielded a channel with preserved channel trafficking and ionic conductance when assayed by two-electrode voltage-clamp (TEVC) in *Xenopus* oocytes (Fig. 1D, SFig. 1A-B). The KCNQ1 HAB linker is an approximately 100 residues long disordered cytosolic loop situated between the two main CaM binding helices in the KCNQ1 CTD ^35,38,39^. Previous studies have found that HAB linker truncation does not affect KCNQ1 function ^50^. The lack of an apparent functional role for the HAB linker may explain the channel’s tolerance to a fluorophore insertion within the HAB linker. Moreover, Cerulean-tagged CaM exhibited higher maximum FRET efficiency with KCNQ1-VenHAB (*E*_Max_∼ 0.34) when compared to labeling Venus at the KCNQ1 carboxy-terminus (*E*_Max_ ∼ 0.17) (SFig. 1 and SFig. 2). This enhanced maximum FRET efficiency is likely due to the closer distance between CaM and the KCNQ1 HAB linker as compared to the KCNQ1 carboxy-terminus, although an orientation-dependent factor cannot be ruled out. We selected KCNQ1-VenHAB as the FRET acceptor in our assay owing to its enhanced maximum FRET efficiency.

To examine binding between CaM and the full-length KCNQ1 channel, we co-transfected Cerulean-tagged CaM and KCNQ1-VenHAB in CHO cells followed by fluorescence imaging using a custom FRET microscope with an EMCCD camera and an optical splitter to simultaneously image fluorescence intensity at the donor and acceptor emission wavelengths (Fig. 1E, see Extended Methods). The acquired images were then analyzed with MATLAB software to compute an apparent FRET efficiency metric *E*D in 1580 cells across 7 independent transfections (Fig. 1E-F, also see Extended Methods). The measured apparent FRET efficiency *E*D observed in each cell depends on two main factors: (1) the true FRET efficiency influenced by the relative orientation and distance between donor and acceptor fluorophores in the bound complex (KCNQ1/CaM in this case), and (2) the binding reaction between the protein pair (KCNQ1 and CaM) that depends on the relative concentrations of the protein pair expressed within the cell (i.e. whether the FRET pairs are bound). Prior studies have shown that the latter factor can be leveraged to quantify relative binding affinity between the labeled FRET pairs. Specifically, variation in transfection efficiency leads to varying titration of free concentrations of FRET donor and acceptor in each cell, enabling fitting of a FRET binding curve by imposing an appropriate binding model ^47,51,52^. Because the KCNQ1/CaM complex exists in a 4:4 stoichiometry with likely identical and independent binding ^35,38^, we fitted the *E*_D_ readouts to a 1:1 (equivalent to 4:4) binding model to derive a FRET binding curve with an apparent dissociation constant (*K*_D,Eff_) related to the binding affinity between CaM and KCNQ1 (Fig. 1F, also see Extended Methods). This analysis yielded a FRET binding curve demonstrating a clear rise in *E*D as the number of free KCNQ1-VenHAB increased with an estimated *K*_D,Eff_ = 30.2 AU (Fig. 1F, blue dots and curve), indicating proximity and binding between Cerulean-tagged CaM and KCNQ1-VenHAB. We note that the estimated *K*_D,Eff_ is in arbitrary units (AU) but can be compared across different constructs for relative binding changes. Although our assay does not report an absolute binding affinity, our method enables quantification of CaM interaction with the full-length KCNQ1 channel within live CHO cells. Furthermore, CaM has been shown to interact with other KCNQ1 regions such as the transmembrane voltage-sensing domain ^10,35,38^, which can be difficult to reconstitute in *in vitro* systems but is fully accounted for in our FRET assay.

To further validate our assay, we measured FRET between Cerulean-tagged CaM and KCNQ1-VenHAB carrying mutations known to disrupt CaM binding to the channel. We mutated the key “IQ” residues in KCNQ1 Helix A (Fig. 1A) known to be important for CaM C-lobe binding ^35,38,39^ to double alanines (I375A/Q376A or IQ/AA). Analysis of FRET results between CaM and KCNQ1-VenHAB-IQ/AA revealed negligible FRET signals beyond those observed with negative controls (Fig. 1F, red, SFig. 1C-D), consistent with the lack of binding between CaM and KCNQ1-IQ/AA. Taken together, our FRET assay detected robust binding between CaM and KCNQ1 WT as well as no interactions between CaM and KCNQ1 IQ/AA designed to ablate CaM binding to KCNQ1. These results validated our assay for probing interactions between CaM variants and the full-length KCNQ1 channel in live cells.

### CaM variants interact with KCNQ1 with different affinities

We next applied our FRET-based assay to probe whether CaM variants bind to the full-length KCNQ1 differently compared to CaM WT. We generated 14 Cerulean-tagged CaM missense variants, including 13 arrhythmia-linked variants (E46K, N54I, N98S, A103V, E105A, G114W, D130G, D132H, D132N, D134H, Q136P, E141G, and F142L) and 1 non-arrhythmia-linked variant identified in the gnomAD database (Q50R) ^17,53^. FRET measurements were performed between each CaM variant and KCNQ1-VenHAB co-transfected in CHO cells under resting or “basal” intracellular [Ca^2+^] conditions. The “basal” intracellular [Ca^2+^] condition was designed to mimic resting low [Ca^2+^] during cardiac diastole and was attained by imaging CHO cells incubated in HBSS solutions containing 1.25 mM Ca^2+^. We then fitted an estimated *K*_D,Eff_ and constructed the 95% confidence interval by bootstrapping for each CaM variants (Fig. 2, SFig. 3, Supplementary Table 1). Statistical significance in binding affinities were determined if the 95% confidence interval of the CaM variant did not overlap with that of the WT control. Our screening identified four CaM variants (E46K, G114W, F142L, and Q50R) which exhibited statistically significant decreased binding affinity to KCNQ1 (increased *K*_D,EFF_) when compared to CaM WT (Fig. 2A,C). The CaM E46K variant featured a 2.6-fold reduction in binding affinity (*K*_D,Eff_ = 70.23 AU E46K vs 30.2 AU WT). Although the reduction in binding is significant, the modest 2.6-fold decrease in affinity suggests that KCNQ1 channels in cardiomyocytes may still pre-associate with CaM E46K depending on the relative concentrations of the CaM E46K variant *vs*. WT. By contrast, the CaM G114W variant exhibited greatly diminished binding to KCNQ1 (Fig. 2A). We were unable to obtain a reliable *K*D,Eff estimate between CaM G114W and KCNQ1 due to the minimal rise in *E*D signals (Fig. 2A), with the best estimate indicating at least 2-orders of magnitude increase in *K*_D,Eff_. Nevertheless, this result indicates KCNQ1 channels in cardiomyocytes carrying the CaM G114W variant are likely devoid of CaM G114W and are instead endowed with CaM WT. Structurally, neither CaM E46 nor G114 are direct Ca^2+^-coordinate residues. CaM E46 is located within the N-lobe EF1-EF2 linker while CaM G114 is situated within the C-lobe EF3-EF4 linker (Fig. 1C). Their positions suggest that CaM E46K and G114W likely disrupt CaM/KCNQ1 binding by altering N-lobe interaction with Helix B and C-lobe interaction with Helix A, respectively. However, potential allosteric effects of each variant on the opposing lobe cannot be ruled out.

**Figure 2.**
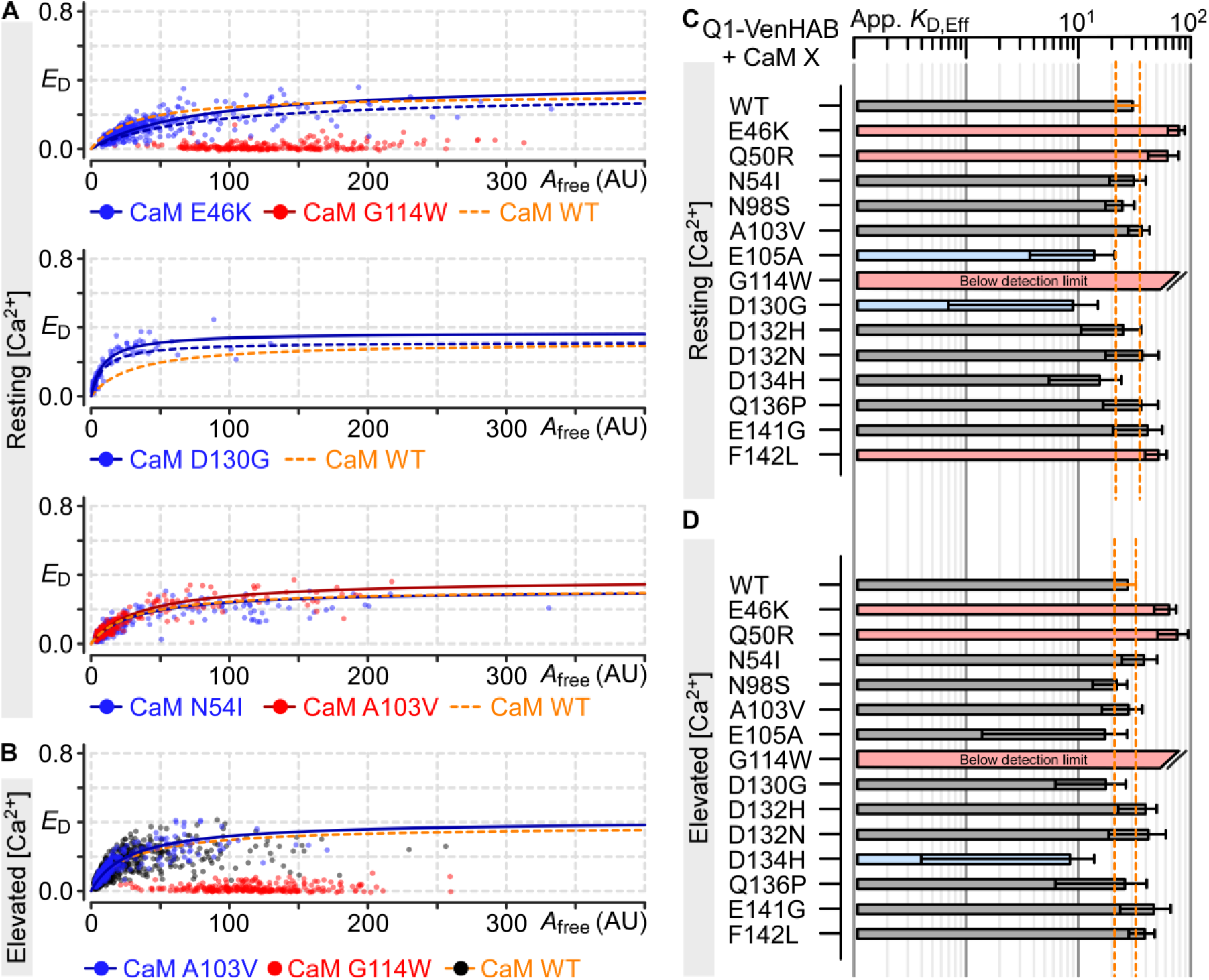
Arrhythmia-associated CaM variants binding to full-length KCNQ1 in live cells. **(A)** FRET binding curves measured between KCNQ1-VenHAB and Cerulean-tagged CaM variants as labeled under resting Ca^2+^ conditions. Each dot is one cell. Dotted blue lines are the FRET binding curves with *E*_Max_ normalized to the same level as WT for *K*_D,Eff_ comparison. Top, middle, and bottom plots illustrate variants exhibiting decreased, increased, and similar binding affinity to KCNQ1-VenHAB, respectively. *A*_free_ corresponds to estimated free concentration of KCNQ1-VenHAB. *n* = 708 (E46K), 291 (G114W), 244 (D130G), 623 (N54I), 686 (A103V). **(B)** Format as in panel A, but showing FRET binding curves measured under elevated Ca^2+^ conditions. *n* = 292 (G114W), 622 (A103V). **(C-D)** Bar plot summary for fitted *K*_D,Eff_ between KCNQ1-VenHAB and the indicated CaM variant under resting or elevated Ca^2+^ conditions. Red and blue bars indicate CaM variants with significantly different *K*_D,Eff_ compared to WT. Error bars are 95% confidence interval. Orange dotted lines denote 95% CI for WT. See Supplementary Tables 1 and 2 for all fitted binding curve parameters.

Majority of variants analyzed (10/14) demonstrated similar or better binding affinity to KCNQ1 when compared to CaM WT under basal intracellular [Ca^2+^] conditions. Two variants (E105A, D130G) exhibited significantly smaller *K*_D,Eff_ estimates indicating enhanced binding affinity to KCNQ1 compared to WT (Fig. 2A,C). The remaining variants such as CaM N54I or CaM A103V yielded similar *K*_D,Eff_ estimates compared to WT (Fig. 2A,C). Taken together, these results demonstrate that most CaM variants can sufficiently compete with their WT counterpart and interact with KCNQ1 in a dominant negative manner. Even among the variants with reduced binding, CaM E46K only showed a mild 2.6-fold reduction. Our screening included variants in multiple CaM regions in both N- and C-lobes, suggesting KCNQ1/CaM interaction is generally tolerant of single missense mutation throughout CaM.

The results so far show that most CaM variants can compete with CaM WT for binding KCNQ1 in a live cell context under basal intracellular [Ca^2+^] conditions. In the cardiac cycle, CaM can coordinate Ca^2+^ ions and change conformation when intracellular [Ca^2+^] rises during systole, thereby potentially changing CaM interaction with KCNQ1. We therefore undertook additional FRET experiments with CaM WT and variants under elevated intracellular [Ca^2+^] conditions to explore potential Ca^2+^-dependent effects. To this end, we incubated transfected CHO cells in solutions containing 10 mM Ca^2+^ and 4 μM ionomycin for at least 15 minutes prior to imaging. This “high” [Ca^2+^] condition was designed to saturate intracellular [Ca^2+^] beyond levels experienced during cardiac systole. FRET binding curves obtained between KCNQ1-VenHAB and cerulean-tagged CaM WT revealed similar binding affinity (*K*_D,Eff_ = 27.3 AU) compared to basal Ca^2+^ conditions (*K*_D,Eff_ = 30.2 AU) (Fig. 2B,D), suggesting that CaM binding affinity to KCNQ1 may not change dramatically between diastole and systole in a cardiomyocyte. Our finding that CaM binding to KCNQ1 is not significantly affected by raising intracellular [Ca^2+^] is consistent with prior studies demonstrating that I_Ks_ ionic current is maximally activated over physiological ranges of intracellular [Ca^2+^] ^40,41^. Next, we performed FRET binding analysis of the CaM variants which generally yielded the same trends as those seen in basal [Ca^2+^] conditions, with most variants exhibiting similar or stronger binding affinity to KCNQ1 (Fig. 2D, SFig. 4, Supplementary Table 2). Among the variants with reduced binding in the basal [Ca^2+^] conditions, the CaM G114W variant maintained a severe loss of interaction with KCNQ1 (Fig. 2B). This finding indicates that the CaM G114W likely does not associate with KCNQ1 at any point during the cardiac cycle. Similarly, the CaM E46K variant featured roughly 2-fold reduced binding affinity (*K*_D,Eff_ = 63.1 AU) compared to WT in high [Ca^2+^] conditions, mirroring the results seen in basal [Ca^2+^] conditions. This suggests that CaM E46K association to KCNQ1 is also likely unaffected by calcium cycling within cardiomyocytes. Curiously, we found one variant, CaM E105A, which showed enhanced affinity to KCNQ1 under basal Ca^2+^ conditions but no significant difference compared to WT under high Ca^2+^ conditions (Fig. 2B,D). However, the confidence interval for CaM E105A was large in our *K*_D,Eff_ estimation, so the results may fall within the margin of error. For most variants, such as CaM A103V, there were no significant differences for CaM variant binding affinity to KCNQ1 under high Ca^2+^ conditions when compared to WT (Fig. 2B, 2D). Taking the combined FRET results together, our findings suggest that most CaM variants feature WT-like binding affinity to KCNQ1 over physiological ranges of intracellular [Ca^2+^]. As these variants are capable of pre-association to KCNQ1, our findings further delineate the CaM variants which may confer KCNQ1 current dysfunction in cardiomyocytes.

### Effect of CaM variants on KCNQ1 membrane trafficking

Given the finding that most CaM variants feature similar binding affinity to KCNQ1 as CaM WT, we next investigated how CaM variants association to KCNQ1 may affect current conduction. Previous studies have shown that CaM facilitates KCNQ1 biogenesis and trafficking to the plasma membrane ^5,6^. Thus, CaM variants may modulate current amplitude by affecting channel membrane trafficking. To explore this possibility, we developed a fluorescence-based assay to compare KCNQ1 membrane trafficking efficiency when co-expressed with CaM variants *vs*. CaM WT. We generated a pseudo-WT (psWT) KCNQ1 construct in which a Cerulean is tagged to the carboxy-terminus and a hemagglutinin (HA) tag is inserted within the extracellular S1-S2 linker between E146 and Q147 (Fig. 3A, KCNQ1-psWT-HA-Cer). The extracellular HA tag enabled targeted fluorescence labeling of KCNQ1 channels on the plasma membrane with an Alexa Fluor 594 (Alexa-594) fluorophore. By contrast, the carboxy terminus-labeled Cerulean afforded an estimation for the total number of KCNQ1 channels within each cell. In each cell transfected with this construct, we defined an apparent trafficking efficiency metric by the fluorescence intensity ratio of Alexa-594 (plasma membrane-bound KCNQ1) to Cerulean (total KCNQ1),

**Figure 3.**
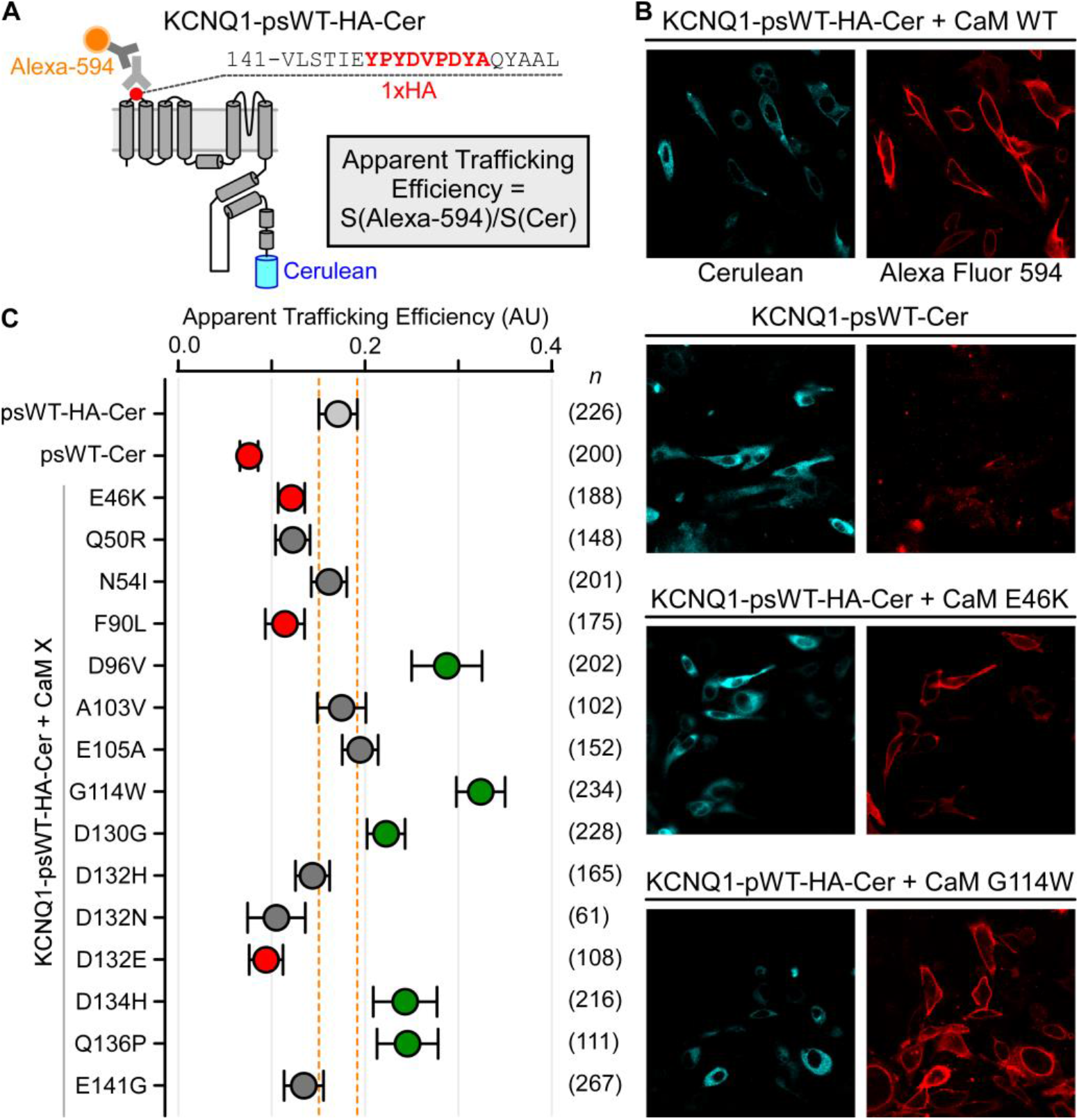
Effect of CaM variants on KCNQ1 membrane trafficking. **(A)** Cartoon illustration of KCNQ1 construct used for membrane trafficking assay (KCNQ1-psWT-HA-Cer). Cerulean fluorophore and HA tag were inserted into KCNQ1 as diagrammed. Total KCNQ1 expressed within the cell correlates with Cerulean signal, while membrane-trafficked KCNQ1 was estimated with Alexa Fluor 594 signal. An apparent trafficking efficiency for each cell was derived by dividing the Alexa Fluor 594 signal to that of Cerulean on confocal microscopy. **(B)** Confocal microscopy images of KCNQ1-psWT-HA-Cer co-expressed with CaM, left cyan and right red images show Cerulean and Alexa Fluor 594 signals, respectively. KCNQ1-psWT-Cer is a construct with a cerulean fused to the carboxy-terminus but no HA tag in the S1-S2 linker. **(C)** Summary data for apparent trafficking efficiency for KCNQ1 co-expressed with different CaM variants. Each dot indicates the mean and error bars represent 95% confidence interval. Orange dashed lines denote 95% confidence interval for WT control. *n* indicates number of cells analyzed for the corresponding CaM variant. Green and red denote statistically significant increased and decreased membrane trafficking when compared to WT control, respectively. Statistical significance calculated with one-way ANOVA followed by Dunnett’s test. See Supplementary Table 3 for all parameters.

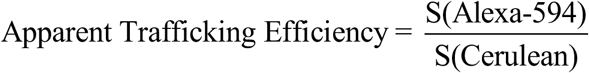

In this assay, we co-transfected CHO cells with KCNQ1-psWT-HA-Cer with CaM WT. 36 hours after transfection, the plasma membrane-bound channels were labeled with a primary anti-HA antibody followed by an Alexa-594 secondary antibody. During staining with the anti-HA antibody, the cells were fixed but not permeabilized through PFA fixation, thus ensuring Alexa-594 were only conjugated on the plasma membrane-bound KCNQ1. The cells were then imaged by confocal microscopy, which revealed robust Cerulean and Alexa-594 fluorescence (Fig. 3B, top 2 panels). Cerulean signal could be seen throughout the cells, while Alexa-594 signals were most evident at the cell membrane, suggesting specific Alexa-594 labeling of membrane-bound channels. Similar fluorescence measurements over 226 cells and revealed an apparent trafficking efficiency of 0.171 ± 0.010 AU (mean ± SEM), establishing a baseline value to which we can compare potential CaM variants effect on KCNQ1 trafficking (Fig. 3C, light gray dot). For negative control, we co-transfected CHO cells with cerulean-tagged KCNQ1 without HA tags in the S1-S2 linker (KCNQ1-psWT-Cer) and performed the same Alexa-594 staining. Confocal imaging of these constructs revealed robust cerulean intensity but minimal Alexa-594 signals (Fig. 3B, second row). The apparent trafficking efficiency computed for this negative control construct over 200 cells was 0.076 ± 0.005 AU (Fig. 3C), demonstrating a clear decrease in our trafficking efficiency metric.

With our assay, we next tested how arrhythmia-associated CaM variants may affect KCNQ1 membrane trafficking by measuring the apparent trafficking efficiency of KCNQ1-HACer co-transfected with various CaM variants (Fig. 3B-C). Significant differences in membrane trafficking were determined by one-way ANOVA followed by Dunnett’s test. We found that seven CaM variants (Q50R, N54I, A103V, E105A, D132H, D132N, E141G) had no effect on KCNQ1 trafficking efficiency compared to CaM WT (Fig. 3C, Supplementary Table 3). As CaM N54I, A103V, D132H, D132N, and E141G exhibited similar binding affinity to KCNQ1 compared to CaM WT (Fig. 2C), the trafficking assay data indicate that these variants likely do not contribute to arrhythmia by modifying I_Ks_ or KCNQ1 current amplitude through changing channel trafficking efficiency.

On the other hand, we found three CaM variants (E46K, F90L, and D132E) that reduced KCNQ1 trafficking efficiency compared to CaM-WT (Fig. 3B-C, Supplementary Table 3). CaM E46K and F90L exhibited mean apparent trafficking efficiency of 0.121 and 0.114 AU, respectively, representing approximately 70% and 67% membrane trafficking compared to WT. These findings suggest that these variants may contribute to arrhythmogenesis in part by reducing I_Ks_ or KCNQ1 current amplitude in cardiomyocytes.

Lastly, we found five CaM variants (D96V, G114W, D130G, D134H, Q136P) increased KCNQ1 trafficking efficiency compared to CaM-WT (Fig. 3B-C, Supplementary Table 3). The variants D134H and Q136P have an apparent trafficking efficiency of 0.243 AU and 0.246 AU or approximately 140% trafficking efficiency compared to WT. As D134H and Q136P bind KCNQ1 with similar affinity as CaM WT, these results suggest that these mutants may contribute to arrhythmogenesis in part by increasing I_Ks_ of KCNQ1 current amplitude. Compared to D134H and Q136P, CaM G114W had a striking apparent trafficking efficiency of 0.324 AU or 180% of WT (Fig. 3B-C). The finding that CaM G114W, which exhibits minimal binding to KCNQ1 (Fig. 2C), increased KCNQ1 trafficking efficiency is interesting. As our assay relies on CaM variant over-expression without knocking down endogenous CaM WT within CHO cells, we interpreted this result as CaM G114W over-expression preferentially binding other CaM targets within each cell. This may increase the fraction of endogenous CaM WT available to bind KCNQ1 within the cell, thereby leading to increased trafficking efficiency. Taken together, these data delineate the arrhythmia-associated CaM variants that affect KCNQ1 membrane trafficking efficiency, which may alter total I_Ks_ or KCNQ1 current amplitude to contribute to arrhythmogenesis.

### Select CaM variants affect baseline KCNQ1 current activation kinetics

In addition to modulating KCNQ1 membrane trafficking, CaM is also known to affect KCNQ1 gating ^5,10^. Previous studies have shown that CaM tunes KCNQ1 function by modulating baseline current activation kinetics through controlling channel entry into the fully activated open state independent of intracellular Ca^2+ 10^. Given the numerous CaM variants exhibiting WT-like interaction with KCNQ1, some of the CaM variants may disrupt KCNQ1 baseline function leading to arrhythmogenesis observed in variant carriers. To examine this possibility, we undertook TEVC experiments in *Xenopus* oocytes. In contrast to the FRET experiments, TEVC was performed with unlabeled KCNQ1 and CaM to avoid potential artifact related to fluorophore fusion. The *Xenopus* oocyte system further enabled direct injection of RNA encoding KCNQ1 and CaM variants, side-stepping potential co-transfection issues in mammalian cell lines. We did not suppress endogenous CaM in *Xenopus* oocyte, which shares 100% amino acid identity with human CaM. This was due to our FRET assay demonstrating most CaM variants can compete with WT, and CaM variants are co-translated with CaM WT in cardiomyocytes *in vivo*. Figure 4A illustrates exemplar ionic current recorded when KCNQ1 was co-expressed with CaM WT and subjected to a series of test voltage pulses. At higher depolarizing pulses, KCNQ1 current featured two distinct time-dependent components of activation (Fig. 4A, red arrow), _fast_ and _slow_. These two components have been previously shown to approximate KCNQ1 entry into distinct open states ^54^. We next screened KCNQ1 co-expressed with CaM variants studied in our FRET assay with TEVC. We replaced the CaM N98S and F142L variants in the TEVC screen with the CaM N98I and D132E variants, as the former two have been previously reported in the context of KCNQ1 ^10,35^.

**Figure 4.**
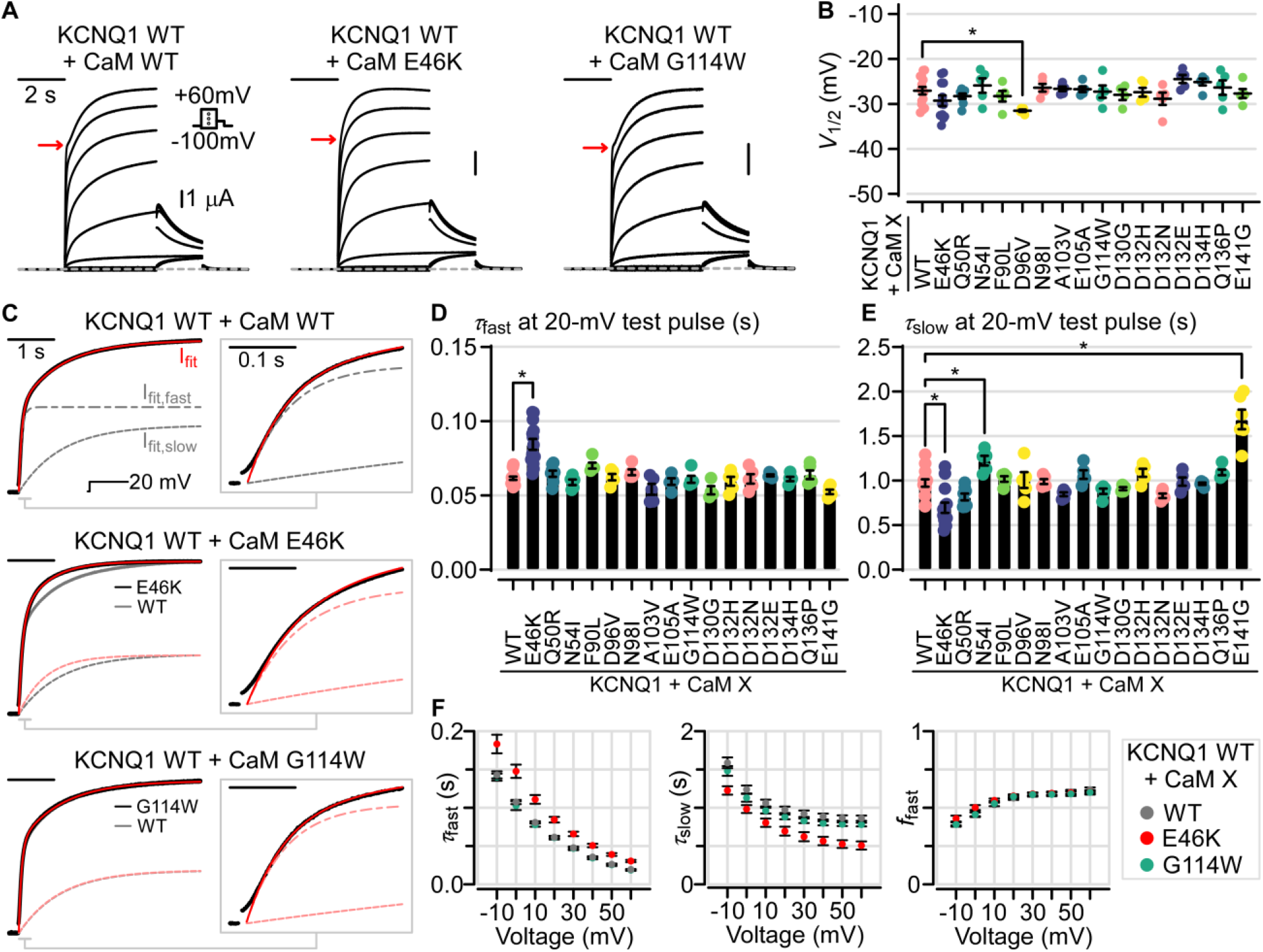
Arrhythmia-associated CaM variants effect on baseline KCNQ1 steady-state activation and activation kinetics. **(A)** Exemplar currents recorded from KCNQ1 WT co-expressed with either CaM WT or CaM variants. **(B)** Summary half-activation voltage (*V*_1/2_) of the conductance-voltage (G-V) curves for KCNQ1 when co-expressed with various CaM variants. **(C)** Exemplar bi-exponential fitting for KCNQ1 activation kinetics. **(D-E)** Summary bar plots for fitted fast and slow time constants for KCNQ1 co-expressed with CaM variants when tested at 20 mV. **(F)** Bi-exponential fitting parameters for KCNQ1 WT co-expressed with CaM WT, E46K, and G114W as a function of voltage. All error bars are SEM.

We first analyzed KCNQ1 steady-state activation property when co-expressed with different CaM variants by fitting the conductance-voltage (GV) relationship for the half-activation voltages (*V*_1/2_), which is the membrane voltage at which KCNQ1 channels are half maximally activated. Statistical significance for all fitted parameters were determined by ANOVA followed by Dunnett’s test. Analysis of measured *V*_1/2_ showed minor differences between WT and all CaM variants probed (Fig. 4B, Supplementary Table 4). CaM D96V demonstrated a subtle but statistically significant 4.4 mV hyperpolarizing shift in *V*_1/2_ (adjusted p-value 0.037). All other variants tested revealed no differences in *V*_1/2_ compared to WT control. These results indicate that most CaM variants do not dramatically perturb the baseline ability of KCNQ1 to open at steady state.

Beyond steady-state response, CaM has been shown to modulate KCNQ1 activation kinetics ^10^. As potassium current activation kinetics figure critically in action potential duration, we next quantified current activation kinetics at 20-mV depolarization by fitting the current tracings to a bi-exponential function with fast and slow components (τ_fast_ and τ_slow_). As shown in Figure 4C, KCNQ1 WT co-expressed with CaM WT yielded ionic currents well-fitted by a bi-exponential function when depolarized to 20 mV (red fit *vs*. black current tracing). We next performed the same bi-exponential fitting procedure on KCNQ1 activation kinetics at 20-mV depolarization when co-expressed with CaM variants (Fig. 4C-E). This analysis revealed that the CaM E46K variant significantly affected KCNQ1 current activation. Specifically, CaM E46K induced a lengthening of τ_fast_ (0.062 WT *vs*. 0.084 E46K, adjusted p-value < 0.0001) and a reduction of τ_slow_ (0.973 WT *vs*. 0.695 E46K, adjusted p-value < 0.0001) when compared to WT (Fig. 4C-E). These two effects together decelerated the fast component and accelerated the slow component, yielding a more rounded appearance for current activation kinetics (Fig. 4A, red arrow). The reduction in τ_slow_ dominated in a 20-mV 4-second test pulse, with KCNQ1 + CaM E46K current reaching steady-state more rapidly than WT as shown by normalized current superposition (Fig. 4C, solid black *vs*. gray). To further test whether E46K affected activation, we performed kinetics fitting at various test voltages and found that it induced consistent increases in τ_fast_ and decreases in τ_slow_ for test voltages ranging from -10 mV to +60 mV (Fig. 4F), providing additional validation for CaM E46K effect on KCNQ1 kinetics. Analysis of other CaM variants revealed that CaM variants N54I and E141G significantly increased τ_slow_ but spared τ_fast_ at 20 mV (Fig. 4E), indicating that CaM N54I and E141G slow KCNQ1 current onset upon membrane depolarization. The ability of the CaM E46K, N54I, and E141G variants to modulate the slow kinetics component is consistent with prior studies demonstrating a role for CaM in modulating the KCNQ1 activation ^10^.

The remaining CaM variants assayed did not yield significant kinetics changes compared to WT at 20 mV (Fig. 4D-E). Notably, the CaM G114W variant induced no changes in either τ_fast_ or τ_slow_ when the test pulse ranged from -10 mV to +60 mV (Fig. 4C-F). As CaM G114W variant features severely reduced binding to KCNQ1, this finding is consistent with the idea that CaM G114W cannot compete with CaM WT for KCNQ1, leading to the channel associating with endogenous CaM WT and exhibiting WT behavior.

Taken together, our results suggest that CaM modulation of KCNQ1 kinetics may be physiologically relevant and further delineate CaM variants that can modulate baseline KCNQ1 current. These kinetic alterations may interact nonlinearly in the cardiac action potential to contribute to arrhythmogenesis.

### Select CaM variants affect baseline I_Ks_ current activation kinetics

Although the KCNQ1-CaM complex constitutes a fully functional voltage-dependent channel, KCNQ1 additionally interacts with the auxiliary subunit KCNE1 in cardiomyocytes to conduct the slow delayed rectifier current (I_Ks_) ^36,37,55^. We will refer to the KCNQ1+KCNE1 complex as I_Ks_. Functionally, KCNE1 causes a significant depolarizing shift in the KCNQ1 steady-state *V*_1/2_ and strongly decelerates current activation kinetics (Fig. 5A), both important factors in the physiological role of I_Ks_. Given the importance of KCNE1 association to KCNQ1 in cardiomyocytes, we undertook additional TEVC analysis to probe how KCNE1 may alter the ability of CaM variants to modulate baseline KCNQ1 current.

**Figure 5.**
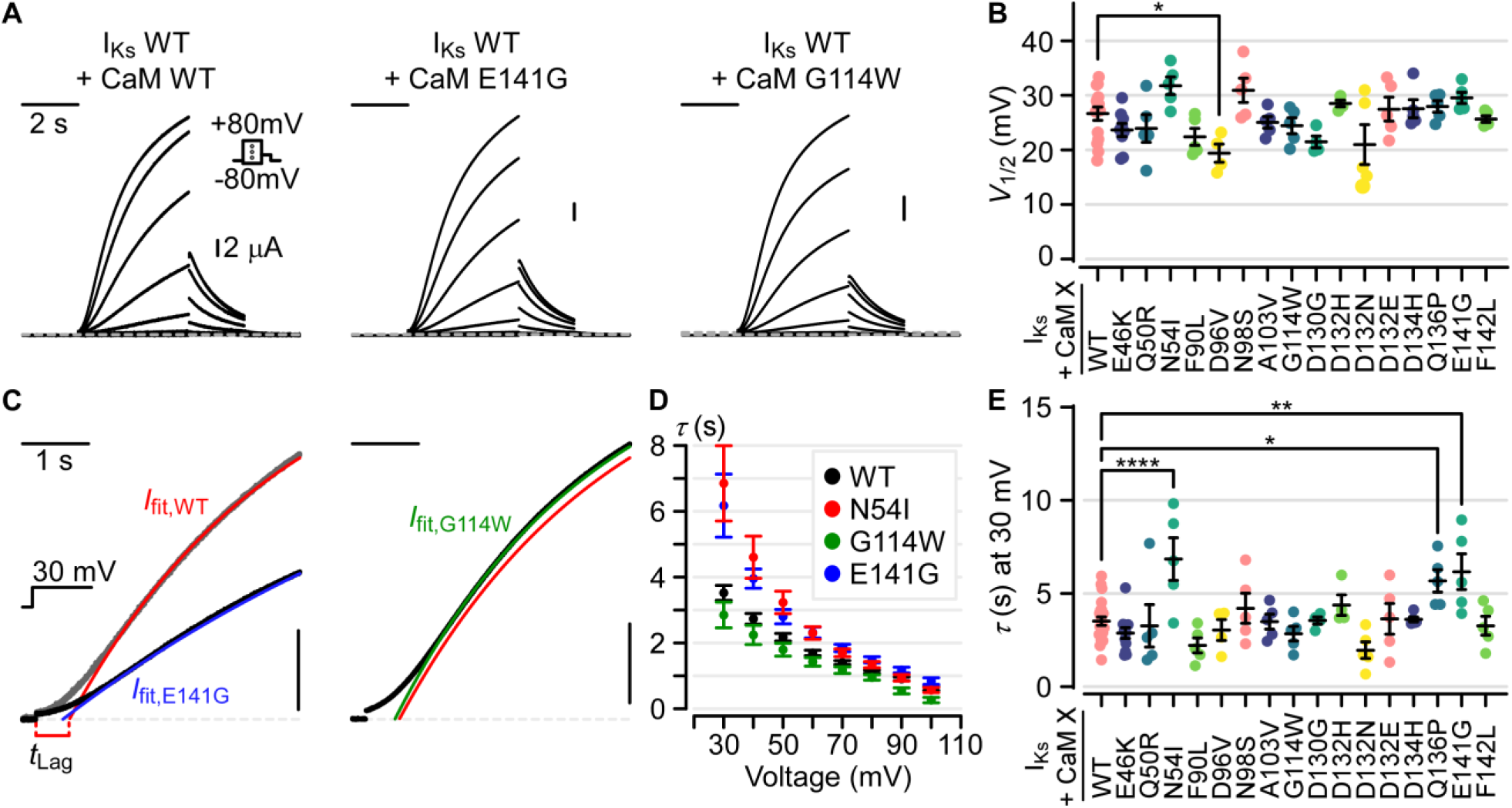
Arrhythmia-associated CaM variants effect on I_Ks_ (KCNQ1+KCNE1) steady-state activation and activation kinetics. **(A)** Exemplar I_Ks_ recordings from *Xenopus* oocytes when co-expressed with CaM WT, E141G, or G114W. **(B)** Summary activation *V*_1/2_ fitted in G-V curves for I_Ks_ co-expressed with CaM variants. Statistical significance determined by one-way ANOVA and Dunnett’s test. **(C)** Exemplar activation kinetics fitting for I_Ks_ co-expressed with CaM variants when depolarized to 30 mV. Currents were fitted to the equation *I*(t) = *A*·exp(−(t - *t*_Lag_) /*τ*), with points prior to *t*_Lag_ excluded from the fitting procedure. Plotted ionic currents were normalized to the fitted *A* for comparison. Vertical scale bars show 25% of normalized current. Left panel: gray and black represent normalized ionic current for I_Ks_ co-expressed with CaM WT and E141G, respectively. Right panel: black and green indicate normalized ionic current and kinetics fit for I_Ks_ co-expressed with CaM E141G. Red line shows identical WT fit seen in the left panel. **(D)** Fitted activation time constant for I_Ks_ co-expressed with CaM WT, N54I, G114W, and E141G as a function of step voltage from 30 mV to 100 mV. Error bars are SEM. **(E)** Summary of fitted activation time constant for I_Ks_ co-expressed with CaM WT or variants when depolarized to 30 mV. Significance was determined by one-way ANOVA followed by Dunnett’s test.

As in the prior KCNQ1 experiments, we first analyzed I_Ks_ steady-state activation when co-expressed with CaM variants in *Xenopus* oocytes. Similar to the results seen in the KCNQ1-only screen, all CaM variants screened exerted minor effects on I_Ks_ baseline half-activation voltage (Fig. 5B, Supplementary Table 5). CaM D96V variant induced a subtle 7.26-mV hyperpolarizing shift in the *V*_1/2_ of I_Ks_ (adjusted p-value 0.034 by one-way ANOVA and Dunnett’s test), mirroring the trend seen in the KCNQ1-only screen. All other CaM variants screened did not show statistically significant differences in I_Ks_ *V*_1/2_ when compared to WT control (Fig. 5B). The comparable effects of CaM variants on KCNQ1 with and without KCNE1 co-expression suggest that KCNE1 does not confer additional ability for these CaM variants to modulate channel opening at steady state.

We next examined whether CaM variants modulate I_Ks_ activation kinetics by fitting the ionic currents with a single exponential function when depolarized to 30 mV. As I_Ks_ activates with a significant delay that cannot be fitted by an exponential function in the early phase, our fits incorporated a time lag (*t*_Lag_) before which the current data were not fitted (Fig. 5C, gray normalized ionic current and red fit) as in prior studies ^56^. Kinetics fitting of I_Ks_ co-expressed with CaM WT revealed an activation time constant (τ) of 3.52 ± 0.24 seconds (mean ± SEM) at 30 mV (Fig. 5C-E). Next, the same fitting procedure was repeated for I_Ks_ co-expressed with CaM variants as summarized in Figure 5E, revealing three CaM variants (N54I, Q136P, and E141G) that significantly prolonged activation kinetics when compared to WT (adjusted p-values < 0.0001, 0.028, 0.0026 for N54I, Q136P, and E141G respectively). CaM E141G increased the activation τ of I_Ks_ to 6.17 ± 0.96 seconds when depolarized to 30 mV, approximately 1.75-fold increase over that of WT and can be appreciated by comparison of normalized currents (Fig. 5C, left panel). Additional kinetics fitting of different step voltages from 30 mV to 100 mV further revealed a consistent increase in τ when I_Ks_ is co-expressed with CaM N54I and E141G when compared to WT (Fig. 5D). Interestingly, CaM N54I and E141G also significantly increased τ_slow_ when co-expressed with KCNQ1 without KCNE1 (Fig. 4E). Taken together, these results suggest that CaM N54I and E141G may slow KCNQ1 and I_Ks_ activation kinetics by a similar mechanism. On the other hand, CaM Q136P did not significantly affect activation kinetics of KCNQ1 without KCNE1 (Fig. 4E) but had an effect on I_Ks_ kinetics (Fig. 5E), suggesting that CaM Q136P requires KCNE1 association to perturb channel gating.

In contrast, the remaining CaM variants did not affect I_Ks_ activation kinetics at 30 mV (Fig. 5E). In particular, CaM G114W did not alter I_Ks_ activation kinetics over voltage ranges from 30 mV to 100 mV (Fig. 5C-D). This result is further consistent with the idea that CaM G114W cannot associate with the channel carboxy-terminus, thus the membrane-trafficked I_Ks_ are endowed with endogenous CaM WT and exhibit WT behavior. Another notable variant is CaM E46K, which induced a decrease in τ_slow_ or hastening of current kinetics in KNCQ1 without KCNE1 (Fig. 4E). Kinetics fitting for I_Ks_ co-expressed with CaM E46K revealed a trend of hastened activation kinetics (mean τ = 2.88 *vs*. 3.52 seconds for CaM E46K *vs*. CaM WT at 30 mV) consistent with the effects seen in KCNQ1 without KCNE1, but ultimately did not reach statistical significance after controlling for multiple comparisons (Fig. 5E). These findings suggest KCNE1 may blunt CaM E46K effect on KCNQ1 activation kinetics, rendering the kinetics effect on I_Ks_ more subtle compared to KCNQ1 alone.

Altogether, our functional screen demonstrates that select CaM variants can modulate baseline I_Ks_ function, with most significant effect on activation kinetics. Moreover, comparison between the I_Ks_ and KCNQ1 results generally demonstrates consistent effects by CaM variants N54I, D96V, and E141G, identifying CaM variants that can affect KCNQ1 gating regardless of KCNE1 association. Our functional screen thus delineates CaM variants (N54I, Q136P, E141G) that can modulate I_Ks_ baseline activation kinetics under resting intracellular Ca^2+^ conditions. As the speed of I_Ks_ activation aids in controlling the cardiac action potential duration, these CaM variants-induced changes in I_Ks_ function may work in concert with effects on other CaM targets to cause arrhythmia.

## Discussion

Missense variants in CaM have emerged in recent years to underlie severe arrhythmia with high mortality rate ^14,16-20^. As CaM targets numerous ion channels important to the cardiac action potential, the mechanisms of CaM-induced arrhythmia (“calmodulinopathy”) are necessarily complex. Multiple studies have shown that CaM variants can significantly impact CaV1.2 and/or RyR2 function ^14,16-18,23,26,27,29,30,32-34,57^. By contrast, although CaM also associates and regulates KCNQ1 channel membrane trafficking and function, relatively little is known regarding whether KCNQ1 or I_Ks_ dysfunction may also contribute to calmodulinopathy. Here, we applied extensive live-cell FRET binding assays, fluorescence-based membrane trafficking assays, and functional electrophysiology to contextualize KCNQ1 within CaM-induced arrhythmogenesis (Fig. 6).

**Figure 6.**
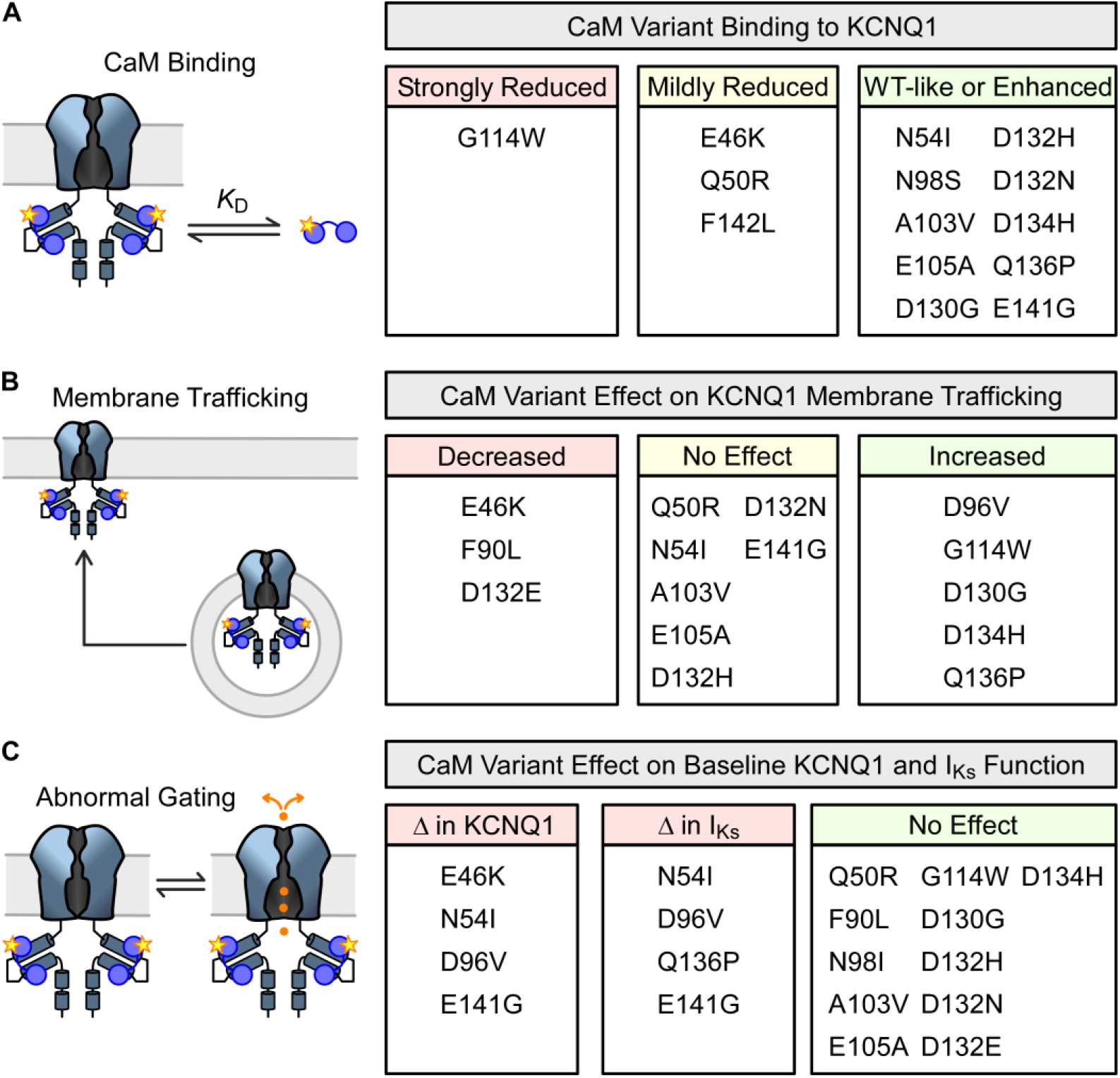
Summary of the effect of CaM variants on KCNQ1. Classification of CaM variants effect on **(A)** KCNQ1 binding, **(B)** KCNQ1 membrane trafficking, and **(C)** KCNQ1 function.

### Most CaM variants can sufficiently compete with CaM WT for binding to KCNQ1

As three independent genes encode CaM within the human genome, CaM variants likely compete with endogenous CaM WT for binding to KCNQ1 within cardiomyocytes. In this study, we examined whether a CaM variant may sufficiently compete with CaM WT with a FRET-based assay that quantify CaM variants binding to KCNQ1 in live cells. As CaM is known to interact with multiple KCNQ1 domains including the voltage-sensing domain, a key strength of our assay is quantification of CaM interaction with the full-length KCNQ1 channel. Our FRET-based screen demonstrates that most CaM variants (10/14 screened) interact with the full-length KCNQ1 with similar or better affinity compared to CaM WT under both resting and elevated intracellular [Ca^2+^] conditions, suggesting that most CaM variants can pre-associate with KCNQ1 in cardiomyocytes expressing both CaM variants and WT (Fig. 2 and Fig. 6A). One potential reason that many CaM variants exhibit similar binding affinity to KCNQ1 compared to WT may be because both CaM N-lobe and C-lobe participate in binding KCNQ1 ^35,38,39,58^. Mutational perturbation of one CaM lobe may be compensated by the other CaM lobe to maintain interaction with KCNQ1. Previous studies have also shown that several CaM variants (e.g. CaM F142L, D96V) bind with similar affinity to other cardiac channels such as CaV1.2 ^29,33^. Our results extend the propensity for CaM variants to exhibit dominant negative interactions to KCNQ1.

Nevertheless, we identified four CaM variants (E46K, Q50R, G114W, and F142L) that exhibit reduced binding affinity to KCNQ1 when compared to WT (Fig. 6A). Three variants (E46K, Q50R, and F142L) feature mildly reduced binding affinity that could feasibly compete with CaM WT for KCNQ1 depending on the relative concentrations of CaM WT to variant within the cell. By contrast, CaM G114W variant represents an interesting case in that the variant appears to be unable to interact with KCNQ1. CaM G114W has been previously shown to exhibit impaired binding to RyR2 CaMBD2 and CaV1.2 IQ domain ^27^, and our result indicates impaired binding extends to KCNQ1. Still, the G114W is significant in the context of KCNQ1 as KCNQ1-CaM interaction generally tolerates single missense mutation in our study. How G114W so strongly impairs CaM interaction with KCNQ1 remains a subject for future studies.

### CaM variants induce aberrant KCNQ1 membrane expression and current conduction

Elucidating CaM variant binding to KCNQ1 represents the first step to contextualize KCNQ1’s contribution to calmodulinopathy, as arrhythmia ultimately arises from cardiac action potential pathologies that are triggered by aberrant ionic currents. In KCNQ1, this may be due to the CaM variant effect on channel membrane trafficking or channel gating. We thus undertook extensive fluorescence and electrophysiology experiments to delineate CaM variant effects on KCNQ1 membrane trafficking and function. As illustrated in Figure 6B-C, our results identified several CaM variants with effect on KCNQ1 membrane trafficking (E46K, F90L, D96V, G114W, D130G, D132E, D134H, Q136P) as well as KCNQ1 or I_Ks_ gating (E46K, N64I, D96V, E141G, Q136P). Given that CaM is well-established to be a key regulator for KCNQ1 membrane trafficking and gating ^5,6,10^, these results highlight KCNQ1 dysfunction as an important consideration when elucidating arrhythmogenesis arising from CaM variants.

As KCNQ1 and I_Ks_ activation kinetics mediate cardiac action potential shortening during β-adrenergic stimulation, how do the observed effects in our study correlate to arrhythmia phenotype? We note that caution should be taken when interpreting how CaM-induced KCNQ1 dysfunction translates to the observed clinical phenotype, as CaM variants can affect multiple channels that non-linearly contribute to the cardiac action potential. Nevertheless, CaM variants with effect on KCNQ1 may be more likely to present with stress or exercise-induced arrhythmia. The CaM E141G and Q136P variants trigger a slower I_Ks_ activation kinetics compared to WT and may provide less repolarizing current, consistent with their association to LQTS ^17^. By comparison, the CaM E46K variant induces a _fast_er KCNQ1 activation kinetics and is associated with CPVT. As CPVT is thought to stem from abnormal Ca^2+^ handling during diastole ^59^, the _fast_er KCNQ1 activation kinetics may contribute to the CPVT phenotype by prolonging diastole. Specifically, KCNQ1 dysfunction caused by CaM E46K may exacerbate the CPVT phenotype during β- adrenergic stimulation where I_Ks_ participates in action potential repolarization. However, CaM E46K is also associated with a reduction in trafficking efficiency, and the activation kinetics effects are more subtle more in I_Ks_ compared to KCNQ1. Adding to the complexity, some variants induce effects seemingly inconsistent with the observed arrhythmia phenotypes. For example, CaM D96V induces a subtle hyperpolarizing shift in KCNQ1 and I_Ks_ steady-state activation as well as increases KCNQ1 trafficking efficiency, both effects expected to increase K^+^ flux and shorten the cardiac action potential. Yet, CaM D96V is associated with a LQTS phenotype ^17^ and a loss of KCNQ1 function would be expected. A likely cause of this difference is that CaM D96V effect on alternate CaM targets such as CaV1.2 may more prominently drive action potential prolongation in cardiomyocytes compared to its effect on KCNQ1. Thus, the KCNQ1 interaction in this case is protective, and loss-of-function KCNQ1 variants in conjunction with D96V may be especially lethal. Lastly, several CaM variants screened in this study feature WT-like binding affinity to KCNQ1 and minimal effects on KCNQ1 and I_Ks_ channel membrane trafficking and gating (Q50R, A103V, E105A, D132H, D132N). Arrhythmia observed in carriers of these CaM variants likely do not arise from direct effect on baseline I_Ks_ or KCNQ1 current.

Clinically, β-blockers are commonly used as medical therapy to treat calmodulinopathy. Still, β-blockers are not equally efficacious in all patients, with more than half of medically treated patients experiencing breakthrough events including sudden cardiac death^17^. Risk stratification for patients most likely to benefit from β-blockade would allow for better therapeutic outcomes. Given the key role of KCNQ1 in the fight-or-flight response, CaM variants that perturb KCNQ1 function (e.g. CaM E141G, Q136P) are more likely to derive therapeutic benefits from β-adrenergic blockade. This study therefore provides data that may inform personalized medical therapy (e.g. β-blocker) in CaM variant carriers.

### CaM variant interaction with KCNQ1 in relation to other CaM targets

In this study, several CaM variants do not alter baseline KCNQ1 gating or membrane trafficking, despite binding KCNQ1 with WT-like affinity. Although these variants likely do not impact the cardiac action potential by directly modulating KCNQ1 current, KCNQ1 may still play a role by altering the variants interaction with other CaM regulatory targets. The pool of free or unbound CaM in cardiomyocytes has been estimated to be severely limited at 50-75 nM representing 1% of total CaM ^60^. The low supply of free CaM may lead to redistribution of CaM variants on distinct CaM targets depending on relative affinities. For example, KCNQ1 may act as a protective “sink” by sequestering CaM variants that bind KCNQ1 with high affinity without affecting I_Ks_ current, effectively chelating these variants from affecting alternate targets.

On the other hand, CaM variants with minimal binding affinity to KCNQ1 (e.g. CaM G114W) may promote CaM G114W to alternate targets and result in surprising findings. CaM binding to KCNQ1 is required for channel assembly and membrane trafficking ^5,6^. Accordingly, CaM G114W is not expected to cause KCNQ1 current dysfunction, as all KCNQ1 expressed on the cardiomyocyte membrane are likely associated with CaM WT. Consistent with this hypothesis, no ionic current abnormalities were detected when KCNQ1 or I_Ks_ were co-expressed with CaM G114W (Fig. 6C). Still, we found that co-expression of KCNQ1 and CaM G114W increased KCNQ1 trafficking efficiency in CHO cells (Fig. 6B and Fig. 3C). We attribute this observation to CaM G114W binding to alternate targets within the cell, thereby freeing more endogenous CaM WT to bind KCNQ1. CaM G114W has been shown to feature moderately reduced, but not abolished, binding to the CaV1.2 and RyR2 as well as impairing CaV1.2 Ca^2+^-dependent inactivation ^27^. It is possible that the lack of binding between KCNQ1 and CaM G114W lead to enhanced CaM G114W pre-association to CaV1.2 and RyR2, thereby exacerbating their dysfunction. Another intriguing possibility is that distinct CaM genes may target their corresponding mRNA to distinct cardiomyocyte cellular regions, with studies suggesting *Calm2* mRNA is more favored to spatially cluster with *Ryr2* mRNA ^61^. Spatial distribution of CaM WT *vs*. variants within cardiomyocytes to distinct CaM targets may add yet another layer of complexity for how CaM variants affect KCNQ1 membrane expression and function. Taken together, the CaM G114W variant highlights the potential complexity of considering distinct CaM targets within the cell, as well as the importance of employing multiple readouts (e.g. binding, membrane trafficking, and functional electrophysiology) to elucidate CaM variant effect.

Taken together, our study furnishes extensive characterization of CaM variants interaction and effect on KCNQ1 channels, delineating the CaM variants that cause KCNQ1 dysfunction to play a role in arrhythmogenesis. This study demonstrates that KCNQ1 dysfunction is a critical component of CaM-induced arrhythmia. The multi-pronged approach employed in this study of measuring binding, surface expression, and channel function can be readily applied to other CaM binding targets. In all, our findings provide key results toward elucidating calmodulinopathy mechanism that integrates numerous CaM regulatory targets.

## Acknowledgements

This work was supported by grants NIH R01 HL136553 and R01 NS092570 (J.R.S.), NIH R01 HL155398 (J.C.), NIH F30 HL151042 (P.W.K.), and US-Israel Binational Science Foundation research grant 2019159 (J.C.).

## Author Contributions

P.W.K., J.C., and J.R.S., conceptualized and designed research; P.W.K., L.W., P.A., J.S., M.M., C.A., D.S., N.S., A.B., performed research and acquired data, P.W.K, L.W., N.S., analyzed data; J.R.S., J.C., funding acquisition; P.W.K., L.W. made figures and wrote original draft; and all authors revised manuscript.

## Data and materials availability

All data required to evaluate the conclusions within this manuscript are presented in the main text and the supplements.

## Materials and Methods

### Molecular biology, cell culture, and transfection

Point mutations were made in KCNQ1 channel and CaM utilizing overlap extension and high-fidelity PCR. DNA sequencing confirmed the presence of all mutants made in the final DNA products. For FRET experiments, CHO cells were cultured in 35-mm dishes and transfected with with jetPRIME or jetOPTIMUS reagents (Polyplus transfection, New York NY) with 4:2 DNA mass-ratio for KCNQ1:CaM corresponding to 0.4:0.2 μg of DNA.

### Electrophysiology and FRET solutions

For all solutions, concentrations are in milli-molar (mM) unless otherwise indicated. *Electrophysiology solutions*. ND96 solution: NaCl 96, KCl 2, CaCl2 1.8, MgCl2 1, HEPES 5, Na pyruvate 2.5, Penicillin-streptomycin 100 U/mL. pH adjusted to 7.6 with NaOH. OR2 Solution: NaCl 82.5, KCl 2.5, MgCl2 1, HEPES 5. pH to 7.6 with NaOH.

### FRET solutions

Hank’s balanced salt solution (HBSS, calcium, magnesium, no phenol red): CaCl2 1.26, MgCl_2_ 0.49, MgSO_4_ 0.41, KCl 5.33, KH_2_PO_4_ 0.44, NaHCO_3_ 4.17, NaCl 137.93, Na_2_HPO_4_ 0.34, D-glucose 5.56. HBSS was purchased from Thermo Fisher Scientific (Waltham, MA). High Ca^2+^ (10Ca^2+^) solution: NaCl 138, KCl 4, MgCl_2_ 1, CaCl_2_ 10, NaHPO_4_ 0.2, HEPES 10, D-glucose 5. pH to 7.4 with NaOH.

### FRET imaging

For all FRET experiments, same-batch transfection of appropriate donor only (e.g. Cerulean-tagged CaM), acceptor only (e.g. Venus-tagged KCNQ1), and spurious FRET constructs (e.g. Venus-tagged KCNQ1 and untagged Cerulean) were also performed and imaged. When FRET data between mutant constructs were collected (e.g. Venus-tagged KCNQ1 and Cerulean-tagged CaM-mutant), the WT FRET pair was also transfected for same-day comparison. Prior to imaging, each plate was washed 3 times with PBS solution and incubated in HBSS(+/+) solution (Thermo Fisher Scientific, Waltham MA) at 37°C for imaging. For experiments involving elevating intracellular Ca^2+^, cells were washed and incubated in 10Ca^2+^ solution with freshly mixed 4 μM ionomycin for at least 15 minutes prior to imaging.

FRET data collection was performed on a Nikon Eclipse Ti-U inverted microscope. Our system design functioned analogously to the “3-cube” FRET technique in which 3 fluorescence channels were acquired ^47,62^, but we implemented multiple bandpass filters, optical splitter, and electronically triggered light source and camera to acquire the 3 channels without using multiple filter cubes. For the Cerulean-Venus or CFP-YFP FRET pair, cells were excited with the Spectra III light engine (Lumencor, Beaverton OR) coupled to the microscope through a liquid light guide with the CFP (integrated bandpass filter: 440/20 nm) and YFP (integrated bandpass filter: 510/25 nm) lines. The microscope was outfitted with a triple-band dichroic beamsplitter (FF459/526/596-Di01, Semrock, Rochester NY) and a triple-band emitter (FF01-475/543/702-25, Semrock, Rochester NY). The microscope’s filter cube allowed both CFP and YFP lines from Spectra III to excite the cells, and simultaneously collected fluorescence emission output from both CFP and YFP at 475 and 543 nm. The fluorescence output was passed into an OptoSplit II Bypass emission image splitter (Cairn Research, Faversham, UK) fitted with a 505nm beamsplitter (T505lpxr, Chroma Color Corporation, McHenry IL), 480nm emission filter (ET480/40m, Chroma Color Corporation, McHenry IL), and 545nm emission filter (ET545/40m, Chroma Color Corporation, McHenry IL). The OptoSplit II Bypass effectively separated the fluorescence output into distinct CFP and YFP emission bands. The two bands were directed toward an Andor iXon EMCCD camera (Andor Technology, Belfast, UK) with each band utilizing one-half of the camera sensor for signal detection driven by the Andor Solis software (Andor Technology, Belfast, UK). The two halves of the camera sensor therefore captured both the CFP and YFP emissions simultaneously. The Spectra III light engine and the Andor iXon EMCCD camera were connected to a Digidata 1440A low-noise data acquisition system (Molecular Devices, San Jose CA) that synchronized both devices by TTL. The Digidata was driven by pClamp software (Molecular Devices, San Jose CA).

During FRET imaging, the camera was used to locate fluorescence cells for imaging through a 20X objective (Nikon CFI S Plan Fluor 20X/0.45). Once an appropriate field of view was found, the Digidata electronically triggered the Spectra III to turn on the CFP and the YFP lines one after the other, during which the Digidata also triggered the camera to acquire two images. The first image corresponded to excitation by the CFP line only, while the second image corresponded to excitation by the YFP line only. As the Optosplit II ensured the camera simultaneously captured CFP and YFP emission, our protocol enabled collection of three fluorescence channels: (1) CFP excitation and CFP emission (donor channel), (2) CFP excitation and YFP emission (FRET channel), and (3) YFP excitation and YFP emission (acceptor channel).

After FRET image acquisition, the images were analyzed and binding curves fitted between KCNQ1 and CaM with MATLAB as detailed in extended methods.

### *Xenopus* oocytes harvesting and two-electrode voltage clamp

*Xenopus laevis* (frogs) were housed in the professional animal facility within the Washington University Danforth campus and oocytes harvested from adult female by laparotomy. All procedures were approved by the Washington University Institutional Animal Care and Use Committee Office. Each oocyte was injected with 10-20 ng of cRNA encoding for KCNQ1, KCNE1, or CaM with a Drummond Nanoject (Broomall, PA). For experiments involving multiple constructs, the cRNA were co-injected at 3:1 (KCNQ1:KCNE1), 1:1 (KCNQ1:CaM), and 3:1:4 (KCNQ1:KCNE1:CaM) mass ratio. Injected oocytes were individually incubated in ND96 in 48-well plates at 18 °C for 2 to 6 days prior to TEVC recording.

### KCNQ1 Membrane Trafficking Efficiency Assay

The trafficking efficiency assay experiments were performed with a Zeiss LSM 880 Airyscan Two-Photon confocal microscope located within the Washington University Center for Cellular Imaging (WUCCI) on the Washington University Medical School campus. CHO cells were transfected in a similar manner to samples prepared for FRET imaging. The cells were plated on 35 mm plates will a small glass-bottom section used for oil immersion imaging. CHO cells were co-transfected with the CaM variant of interest and KCNQ1-psWT-HA-Cer (Fig. 3A). In each experiment, the Alexa-Fluor 594 were conjugated to the plasma membrane-bound KCNQ1 via a secondary antibody targeted to the HA tag within the S1-S2 linker. During staining, cells were fixed but not permeabilized through PFA fixation to minimize intra-cellular labeling. To prevent non-specific labeling of Alexa-594, cells were incubated overnight with BSA at 4 °C after fixation. As all KCNQ1 subunits are labeled with a Cerulean probe, the fluorescence intensity in the Cerulean channel is proportional to the total number of KCNQ1 in each cell. On the other hand, the fluorescence intensity of the Alexa-594 channel is proportional to the number of plasma membrane-bound KCNQ1, as the HA-tag labeling site is within the extracellular S1-S2 linker. The apparent trafficking efficiency of KCNQ1 was computed for each cell with the equation

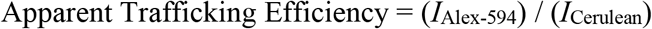

Where *I*_Alex-594_ and *I*_Cerulean_ are the fluorescence intensities of the Alexa-694 and Cerulean channels, respectively. All measurements were taken at the same PMT and laser intensity levels. Statistical results are ANOVA tests compared to the WT CaM trafficking efficiency.

### Electrophysiology data analysis

Data analysis performed with MATLAB (MathWorks, MA). Conductance–voltage (*GV*) curves were quantified by fitting a single Boltzmann function. The instantaneous tail currents following test pulses were normalized to the maximum tail current and fitted with the equation

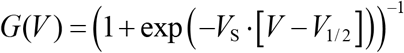

where *V* is the test voltage, *V*S is the slope factor controlling the steepness of the Boltzmann equation, and *V*_1/2_ is the half-activation voltage. *V*_S_ is further related to *RT*/*zF*, where *R* is the gas constant, *T* is the absolute temperature, *z* is the equivalent valence, and *F* is the Faraday constant.

KCNQ1 current activation kinetics were quantified by fitting the 4-sec test-pulse currents with a biexponential function in the general form of

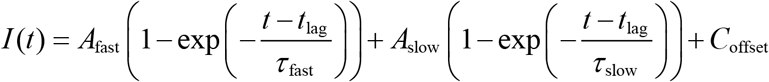

with one fast component and one slow component. Each exponential is characterized by a time constant τ as well as a steady-state amplitude *A*. The variable *t*_lag_ corresponds to the time lag to begin kinetics fitting. Current data between the start of the test pulse and *t*_lag_ (i.e. I(0 < *t* < *t*_lag_)) were not included in the fit. Kinetics fitting of I_Ks_ current activation used a similar equation, but only a single exponential component was used.

For fitting, each current trace was baseline corrected with the mean current at the holding potential (−80 mV) for each trace. With KCNQ1 specifically, as the slow component contained more data points than the fast component, the fast current component was first estimated by fitting the first 0.5 second after the test pulse. A second overall fit was then applied to the entire 4-second test pulse current, with the fast time constant constrained to ±25% of the first fit. For KCNQ1 current activation kinetics, the fraction of the total current carried by the fast component was calculated with the following equation

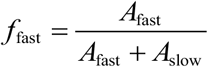

Because KCNQ1 current activates without very significant lag, *t*_lag_ variable was set to zero. On the other hand, *t*_lag_ was a free parameter in I_Ks_ kinetics fitting. *t*_lag_ was determined automatically by linear extrapolation. First, the derivative of the current trace was estimated and smoothed by the “diff” and “smooth” functions on MATLAB, respectively. The value of the maximum rate of change (*m*_rate,max_) and the time point at which maximum rate of change occurred (*t*_rate,max_) within the first 0.4 second was then determined. Using these two parameters, a tangent line was drawn on the current trace at *t* = *t*_rate,max_ with slope *m*_rate,max_. Finally, *t*_lag_ was determined by linear extrapolating the tangent line back to the time point where the line intersects with the instantaneous current at beginning of the test pulse using the equation,

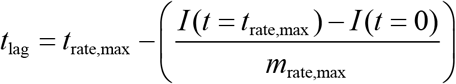

